# Brown Adipose Tissue Remodeling Precedes Cardiometabolic Abnormalities Independent of Overweight in Fructose-Feed Mice

**DOI:** 10.1101/615674

**Authors:** Thaissa Queiroz Machado, Debora Cristina Pereira-Silva, Leidyanne Ferreira Gonçalves, Caroline Fernandes-Santos

**Author notes:** Both authors contributed equally to this work (Thaissa Queiroz Machado, Debora Cristina Pereira-Silva). Corresponding Author: Rua Dr. Silvio Henrique Braune, 22, Centro, Nova Friburgo, RJ, Brazil, 28.625-650.

## Abstract

**Objectives:** To investigate the early cardiometabolic abnormalities along with WAT and BAT remodeling in short-term fructose feeding mice model.

**Methods:** Mice (n=10-11/group) were fed for four weeks with control diet (AIN93-M) or experimental diets rich in glucose or fructose. We investigated body weight, body adiposity, blood glucose, lipid and hepatic parameters, and white (WAT) and brown adipose tissue (BAT) histopathology.

**Results:** Fructose feeding promoted neither weight gain nor hypertrophy of visceral and subcutaneous WAT depots, but the fat was redistributed toward visceral depots. Glucose, lipid and hepatic metabolic dysfunction were not yet noticed in fructose-fed mice, with the exception for an elevation in total cholesterol and hepatic weight without steatosis. BAT mass did not increase, and it was proportionally reduced compared with visceral WAT in fructose feed mice. BAT suffered premature adverse morphological remodeling, characterized by increased lipid deposition per tissue area in enlarged intracellular lipid droplets.

**Conclusion:** Short-term fructose feeding redistributes body fat, changes the proportion of BAT to visceral fat, and promotes BAT adverse remodeling, characterized by enlarged intracellular lipid droplets.

## Introduction

Overfat is strongly associated with insulin resistance and chronic inflammation, as well as hypertension, dyslipidemia, cardiovascular diseases (for instance, coronary heart disease and stroke), cancer, type 2 diabetes, gallbladder disease, osteoarthritis, gout, and sleep apnea (“Obesity: Preventing and Managing the Global Epidemic. Report of a Who Consultation” 2000). However, excess body fat not always appears to be associated with cardiometabolic abnormalities. This statement came from observations that some subjects have normal weight but display excessive body fat percent and cardiometabolic dysfunction (metabolically obese normal weight, MONW), and there are also metabolically healthy but obese (MHO) subjects, that are overfat but have no signs of cardiometabolic dysfunction (Ruderman, et al. 1998). Regardless of body weight and body mass index, excessive body fat is associated with cardiometabolic dysfunction. The earliest signs of cardiometabolic dysfunction are excessive body fat, insulin resistance, and chronic low-grade systemic inflammation (Maffetone, Rivera-Dominguez, and Laursen 2017).

Individuals without obesity but with dyslipidemia and metabolic abnormalities have also been termed as normal weight dyslipidemia (NWD) (Ipsen, Tveden-Nyborg, and Lykkesfeldt 2016). This population has an increased risk of developing nonalcoholic fatty liver disease (NAFLD), cardiovascular disease and type 2 diabetes (Pagadala and McCullough 2012; Ruderman, et al. 1998). As the liver regulates lipid metabolism and plasma lipid levels, it plays a significant role in turning an unhealthy diet and lifestyle into an unbalanced metabolic profile. The product of fasting triglycerides and glucose (TyG) is a useful index to assess both insulin resistance and hepatic steatosis/nonalcoholic steatohepatitis (NASH) in apparently normal subjects (Simental-Mendía, et al. 2016; Simental-Mendía, Rodríguez-Morán, and Guerrero-Romero 2008). Thus, it could be used as an additional early marker of metabolic dysfunction in MONW individuals.

The liver and the white adipose tissue (WAT) continuously exchange very low-density lipoproteins (VLDL) and free fatty acids (FFA) to store (WAT) and distribute (liver) energy. In obesity, the liver modulates WAT inflammation and insulin sensitivity, and the hypertrophic WAT influences liver metabolism and inflammation (Scheja and Heeren 2016). Although WAT is a specialized lipid storage organ for excess calories, the brown adipose tissue (BAT) contains many mitochondria to dissipate chemical energy. It has been proven that BAT activity controls plasma clearance of circulating triglyceride rich lipoproteins (TRL) by increasing its uptake into BAT and thus promoting its turnover (Bartelt, et al. 2011).

So far, an animal model that mimics the MONW phenotype is missing. The Goto-Kakizaki rat is a non-obese Wistar substrain which develops type 2 diabetes early in life, characterized by mild hyperglycemia, insulin resistance, hyperinsulinemia and mild inflammation. It was proposed as a MONW model (Denis and Obin 2013), but it is not accessible to all researchers worldwide. A potential MONW model is fructose overfeeding in rodents. Purified diets containing fructose are capable of elevating TG and hepatic glucose production, ultimately leading to insulin resistance and hypertrygliceridemia (R. and A. 2017). The C57BL/6 mice feed with fructose for 8 weeks have no increase in body weight, but displays increased adiposity, glucose intolerance, and ectopic hepatic lipid accumulation as TG and diacylglycerol (Montgomery, et al. 2015).

Overall, the role of BAT on MONW phenotype was not investigated so far, and we hypothesize that BAT changes run together with cardiometabolic abnormalities independent of body weight gain. Thus, the aim of the present study was to investigate the early cardiometabolic changes along with WAT and BAT remodeling in C57BL/6 mice by short-term fructose feeding.

## Material and Methods

### Experimental design

The handling and experimental protocols were approved by the local Ethics Committee to Care and Use of Laboratory Animals (CEUA#647/15). The study was performed in accordance with the Animal Research Reporting in Vivo Experiments ARRIVE guidelines and the Guideline for the Care and Use of Laboratory Animals (US NIH Publication N° 85-23. Revised 1996) (Kilkenny, et al. 2010). Male C57BL/6 mice at two months of age were obtained from colonies maintained at the Federal Fluminense University and kept under standard conditions (12 h light/dark cycles, 21±2°C, humidity 60±10% and air exhaustion cycle 15 min/h).

At three months old, mice were randomly allocated into three groups according to the diet offered (n=10-11/group). Control group received a purified diet according to AIN-93M standards (Reeves, Nielsen, and Fahey 1993), and the other two groups received isoenergetic modified AIN-93M diets rich in glucose or fructose (3.81 kcal/g), acquired from Pragsolucoes (Jau, Sao Paulo, Brazil). Both glucose and fructose diets were rich in simple carbohydrates since they have lower complex carbohydrate content (corn starch) and sucrose was removed (Table 1). Glucose rich diet was administered to evaluate if changes encountered by fructose feeding were due solely to single carbohydrate overfeeding or to the quality of the carbohydrate. Food and water were offered *ad libitum*. Food intake was measured daily and body mass weekly throughout 4 weeks. Energy efficiency was calculated as [(Δ body weight/ ∑ Kcal ingested) ×100].

**Table 1.** – Diets composition

| Ingredients (g/Kg) | Diets |  |  |
| --- | --- | --- | --- |
|  | Control | Glucose | Fructose |
| Casein | 140.0 | 140.0 | 140.0 |
| Cornstarch | 620.7 | 220.7 | 220.7 |
| Sucrose | 100.0 | - | - |
| Fructose | - | - | 500.0 |
| Glucose | - | 500.0 | - |
| Fiber | 50.0 | 50.0 | 50.0 |
| Soybean oil | 40.0 | 40.0 | 40.0 |
| Vitamin mix* | 10.0 | 10.0 | 10.0 |
| Mineral mix* | 35.0 | 35.0 | 35.0 |
| Choline bitartrate | 2.5 | 2.5 | 2.5 |
| L-Cystine | 1.8 | 1.8 | 1.8 |
| Antioxidant<sup>#</sup> | 0.008 | 0.008 | 0.008 |
| Sum, g | 1 000.00 | 1 000.00 | 1 000.00 |
| Kcal/g | 3.81 | 3.81 | 3.81 |

\* Vitamin and mineral mix composition is based on AIN-93M.
<sup>#</sup> Tert-butylhydroquinone.

**Table 2.** – Energy intake, body weight and adiposity

| Parameters | Groups |  |  |
| --- | --- | --- | --- |
|  | Control | Glucose | Fructose |
| Ingestion |  |  |  |
| Σ FI, g | 115.86±1.92 | 116±1.85 | 103.79±2.71<sup>b</sup> |
| EE, mg/Kcal | 5.97±0.88 | 2.20±0.68<sup>a</sup> | 0.76±0.36<sup>b</sup> |
| Body weight |  |  |  |
| Initial, g | 30.36±0.46 | 30.99±0.61 | 31.36±0.46 |
| Final, g | 32.99±0.29 | 31.96±0.52 | 31.68±0.54 |
| Δ, g | 2.63±0.39 | 0.97±0.31<sup>a</sup> | 0.32±0.15<sup>b</sup> |
| WAT |  |  |  |
| Genital, mg/g of BW | 12.65±0.82 | 12.05±0.78 | 14.04±0.46 |
| Retroperitoneal, mg/g of BW | 3.78±0.30 | 3.29±0.22 | 4.10±0.32 |
| Inguinal, mg/g of BW | 11.51±0.68 | 10.88±0.83 | 11.38±0.66 |
| Visc:Subc ratio | 1.44±0.06 | 1.43±0.06 | 1.65±0.05<sup>a,b</sup> |
| Adipocyte diameter |  |  |  |
| Genital, μm | 59.43±1.69 | 56.43±1.84 | 57.65±0.96 |
| Retroperitoneal, μm | 54.03±0.73 | 52.99±3.02 | 54.59±1.50 |
| Inguinal, μm | 42.27±2.51 | 37.79±2.41 | 35.68±1.98 |

Data are expressed as mean±SEM. When indicated, P<0.05, [a] ≠ Control group and [b] ≠ Glucose group (one-way Analysis of variance and Tukey's multiple comparison test). Abbreviations: Σ FI, sum of food intake, from day 0 to the 30<sup>th</sup> day; EE, energy efficiency; WAT, white adipose tissue; BW, body weight; Visc, visceral fat; Sub, subcutaneous fat. Energy efficiency was calculated based on Δ body weight and Σ energy intake.

### Glucose, lipid and hepatic parameters

On the day of euthanasia, blood was obtained from awake 6-hour fasted mice by milking the tail after a little incision on its tip and plasma glucose was assessed using a glucometer (One Touch Ultra, Johnson& Johnson, SP, Brazil). Mice were then deeply anesthetized with ketamine 100.0 mg/kg (Francotar®, Virbac, Brazil) and xylazine 10.0 mg/kg ip (Virbaxyl 2%®, Virbac, Brazil), and the heart was exposed for blood collection (right atrium). Blood was allowed to clot, centrifuged (1,500 × g) and the serum was stored at −80°C for total cholesterol, HDL and triglyceride (TG) colorimetric assay (cat#K083, #K071and K117 respectively, Bioclin, Quibasa, Belo Horizonte, Minas Gerais, BR) and insulin Elisa assay (cat#EZRMI-13K, Merck Millipore, Billerica, MA, EUA) according to manufacturer’s instructions. Insulin resistance was evaluated by the homeostatic model assessment, where HOMA-IR = [insulin (µU/mL) × glucose (mmol/L)]/22.5 (Matthews, et al. 1985) and hepatic TG content was determined as described elsewhere (Gonçalves, et al. 2017). TyG index, the product of fasting plasma glucose (FPG) and TG was used to further assess insulin resistance and liver steatosis (Simental-Mendía, Rodríguez-Morán, and Guerrero-Romero 2008; Simental-Mendía, et al. 2016). It was calculated as ln[FPG (mg/dL) × TG (mg/dL)/2] (Simental-Mendía, Rodríguez-Morán, and Guerrero-Romero 2008).

### Fat harvesting

Interscapular brown fat and white visceral (perigonadal and retroperitoneal) and subcutaneous (inguinal) fat pads were carefully dissected from both sides of the animal, weighed and then immersed in 4% phosphate buffered formalin pH 7.2 for 48 h. Samples of both contralateral fat pads were submitted to routine histological processing, embedded in paraplast, sectioned 3 μm thick and stained with hematoxylin and eosin. To calculate fat distribution, WAT was considered as ∑ (perigonadal (mg) + retroperitoneal (mg) + inguinal (mg)]), visceral WAT as ∑ (perigonadal (mg) + retroperitoneal (mg)]) and the inguinal depot as subcutaneous WAT.

### Adipocyte morphometry

Digital images were obtained from histological sections using a Leica DMRBE microscope (Wetzlar, German) coupled to a video camera Kappa (Gleichen, German). Morphometry was performed in the Image-Pro® Plus software v. 5.0 (Media Cybernetics, Silver Spring, MD, USA). In the interscapular BAT, eight nonconsecutive images were acquired to assess brown adipocyte diameter, lipid droplet (LD) diameter, and the percentage of tissue area occupied by LD. For lipid area, a selection tool was used to mark the pixels that represented lipid droplets. The selection was segmented in a new digital image in black and white, where the white color represented the LD, and the black color represented the remaining tissue. Then, the area occupied by the white color was quantified through the image histogram tool. In the three WAT depots studied, we assessed adipocyte diameter by measuring their smallest and largest diameters, as previously described (Fernandes-Santos, et al. 2009). In this case, we used six animals per group, four nonconsecutive images per animal, and randomly measured 10 adipocytes per image, totalizing 40 adipocytes per mice.

### Statistics

Data are expressed as mean ± SEM and tests to assess normality and homoscedasticity of variances were run. Comparison among groups was made by ANOVA one-way followed by a post-hoc test of Tukey. A *P*-value of 0.05 was considered statistically significant (GraphPad® Prism software v. 6.0, La Jolla, CA, USA).

## Results

### Early signs of fructose overfeeding: Body fat redistribution without body weight gain

Despite glucose and fructose diets were isoenergetic compared to the control diet, Table 1 shows that cumulative food intake (FI) in fructose group was 10% lower than the control group (*P*<0.02). Additionally, energy efficiency reduced by 63% and 87%, respectively, in glucose and fructose groups (*P*<0.0001). Neither glucose nor fructose feeding changed body weight, and surprisingly BW gain during the 4 weeks on experimental diets was slowed down, since Δ BW was 63% and 88% lower in glucose and fructose groups, respectively, compared to control group (*P*<0.0001). All white fat depots studied did not vary in weight and did not present adipocyte hypertrophy after glucose or fructose feeding. However, fructose feeding changed body fat distribution, since the ratio between visceral and subcutaneous white fat increased, compared to control group (+15%, *P*=0.016).

### Short-term fructose feeding had limited impact on glucose, lipid and hepatic metabolism

Thirty days of glucose or fructose feeding did not modulate glucose and lipid metabolism to a great extent, as shown in Table 3. Glucose group presented a decrese of 41% on blood glucose (*P*=0.0002) that was followed by an increase in serum (+80%, *P*<0.0001) and liver (+79%, *P*=0.0002) triglyceride. On the other hand, fructose only affected total cholesterol (+32%, *P*=0.002). Insulin resistance, assessed by HOMA-IR and TyG index, was not developed after 4 weeks of glucose or fructose feeding. Although fructose feeding increased liver weight, TyG index did not point to the presence of hepatic steatosis (Table 3), and we also did not find histopathological changes compatible with steatosis (data not shown).

**Table 3.** – Glucose, lipid and hepatic metabolism

| Parameters | Groups |  |  |
| --- | --- | --- | --- |
|  | Control | Glucose | Fructose |
| Glucose metabolism |  |  |  |
| Glucose (day 0), mg/dL | 128±7.11 | 116.9±7.04 | 107±4.31 |
| Glucose (day 30<sup>th</sup>), mg/dL | 120.6±8.56 | 68.6±5.45<sup>*a</sup> | 111±5.67<sup>b</sup> |
| Insulin, ng/mL | 0.32±0.01 | 0.29±0.02 | 0.38±0.01 |
| HOMA-IR | 2.66±0.24 | 2.44±0.25 | 2.93±0.18 |
| Lipid Metabolism |  |  |  |
| Total Cholesterol, mg/dL | 92.64±4.34 | 95.61±3.11 | 122.1±5.81<sup>a,b</sup> |
| HDL, md/dL | 47.16±2.04 | 49.11±3.13 | 41.2±4.51 |
| Triglyceride, mg/dL | 71.49±4.18 | 128.7±15.51<sup>a</sup> | 65.92±3.62<sup>b</sup> |
| Hepatic metabolism |  |  |  |
| Liver, g/g BW | 0.042±0.0006 | 0.041±0.0007 | 0.044±0.0006<sup>a</sup> |
| Triglyceride, mg/dL/mg | 2.76±0.34 | 4.95±0.54<sup>a</sup> | 2.78±0.20<sup>b</sup> |
| TyG index | 8.37±0.13 | 8.31±0.07 | 8.11±0.09 |

Data are expressed as mean±SEM. When indicated, P<0.05, [a] ≠ control group, [b] ≠ glucose group (one-way Analysis of variance and Tukey's multiple comparison test), and [\*] ≠ glucose group of blood glucose (day 0, paired t Test). Abbreviations: BW, body weight; TyG, product of glucose and triglyceride.

### Early changes in BAT morphology due to short-term fructose feeding

Figure 1 shows that BAT weight remained unchanged after 4 weeks of glucose or fructose feeding. The proportion between BAT and WAT was also not changed, despite a subtle decrease in fructose group. When the amount of visceral WAT is compared to BAT mass, there is a significant decrease of 18% in fructose group compared to control group (*P*=0.03). Figure 2 depicts BAT morphological remodeling. Thirty days of glucose or fructose feeding did not change brown adipocyte size (Fig 2A), but intracellular lipid deposition is already noticed by glucose and fructose overfeeding (Fig 2B). The percentage of BAT area occupied by LD increased 19% (*P*=0.01) in glucose group and 17% (*P*=0.02) in fructose group. In fructose group, this increase can be attributed to LD hypertrophy, since average LD diameter increased 17% (*P*=0.006) compared to control group (Fig 2C). BAT photomicrographs in Fig 2 D-I show that lipid droplets have a uniform size and are evenly distributed in control mice (D, G). Although glucose group did not present lipid droplet hypertrophy (Fig 2B), some large droplets are noticed in Fig 2H. Finally, fructose feeding lead to lipid droplet hypertrophy as seem in Fig 2E-F.

**Figure 1.**
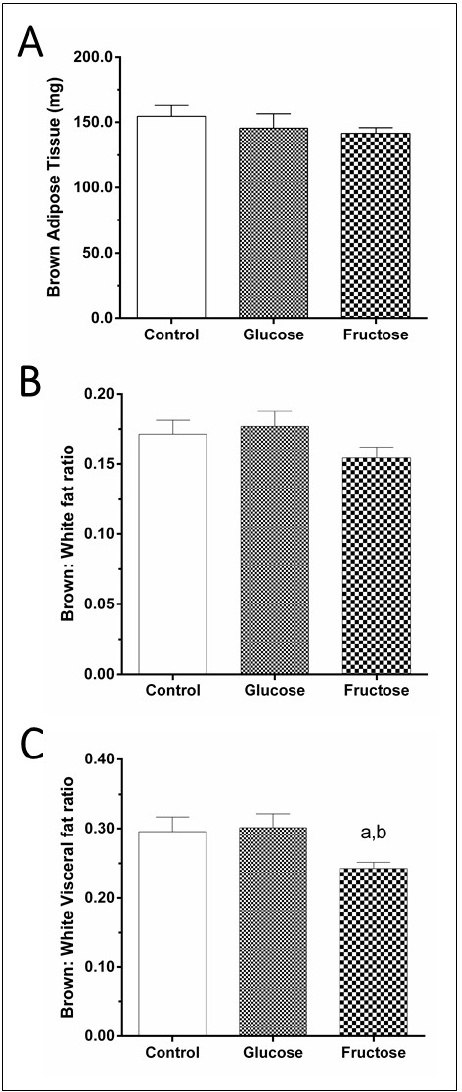
Interscapular brown fat. Glucose or fructose feeding neither change brown adipose tissue (BAT) mass (**A**) nor the ratio between BAT and white adipose tissue (WAT) mass (**B**). However, fructose decreased the ratio between BAT and visceral WAT (**C**). N=6 mice/group, mean ± SEM, One-way analysis of variance, post-hoc test of Tukey, P<0.05, [a] vs. control group and, [b] vs. glucose group.

**Figure 2.**
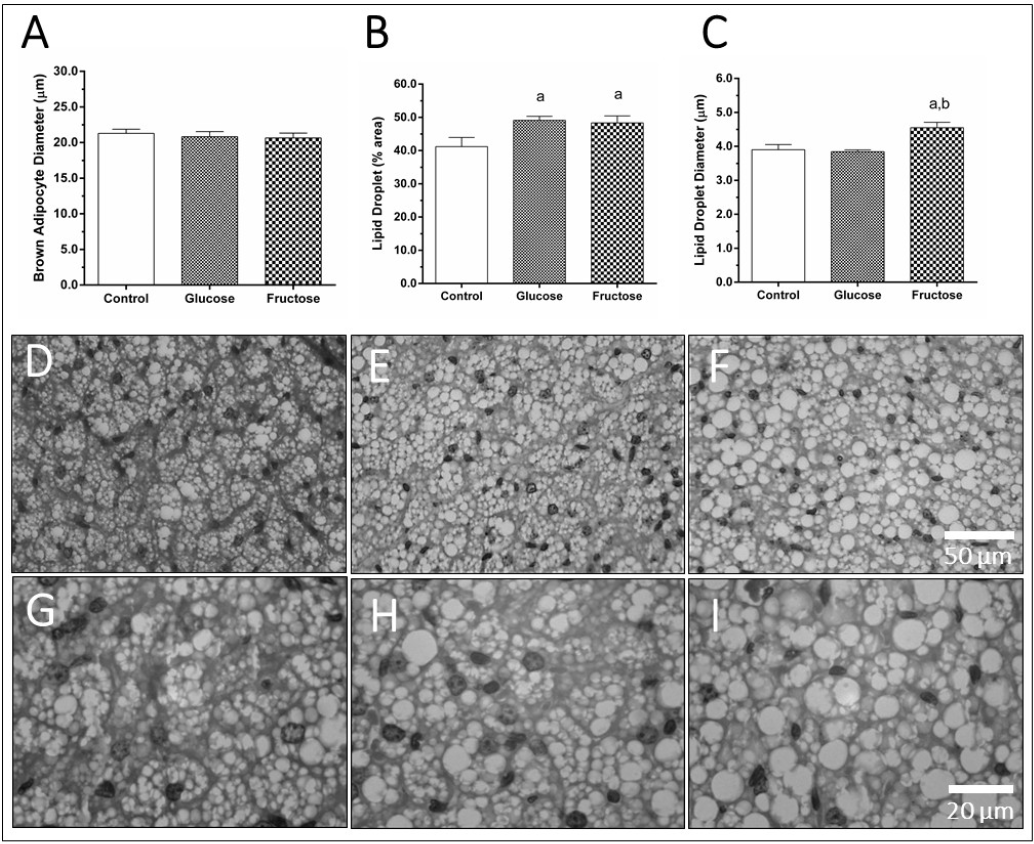
Diet-induced morphological changes in interscapular brown adipose tissue (BAT) of male C57BL/6 mice. Brown adipocytes size was not altered by glucose or fructose feeding (**A**), but both diets increased the percentage of tissue area occupied by cytoplasmic lipid droplets (**B**). In fructose-feed mice, the last change was due to an increase in lipid droplet diameter (**C**). D-I are photomicrographs (H&E stain) in lower (**D**, **E**, **F**) and higher (**G**, **H**, **I**) magnification illustrating BAT remodeling. Lipid droplets are uniformly distributed in size along the tissue in control mice (**D**, **G**). Although glucose feeding did not alter average lipid droplet size, some large droplets are noticed (**E**, **H**). On the other hand, lipid droplet hypertrophy is visible in **E** and **F** due to fructose feeding. N=6 mice/group, mean ± SEM, One-way analysis of variance, post-hoc test of Tukey, P<0.05, [a] vs. control group and, [b] vs. glucose group.

## Discussion

We demonstrated in male C57BL/6 mice that short-term fructose feeding did not promote weight gain and adiposity, but WAT fat was redistributed toward visceral depots. Glucose, lipid and hepatic metabolic dysfunction were not yet noticed, with the exception of increased serum total cholesterol and liver weight. The most prominent finding is that fructose feeding changes the proportion between BAT and visceral WAT mass, where the second predominates, and BAT suffers an important premature morphological remodeling due to increased lipid deposition in enlarged intracellular LD, which precedes the onset cardiometabolic abnormalities.

In the present study, fructose diminished cumulative food intake, and it likely led to an absence of body weight gain. Tillman et al showed that even when cumulative food intake is increased, no significant increment in final body weight is seen after 14 weeks of 60% fructose feeding (Tillman, et al. 2014). They also showed that metabolic rate is increased in the second and ninth weeks of fructose feeding, but at the fourteenth week metabolic rate is similar to control group. We showed through energy efficiency that fructose group gained less weight per energy consumed, compared to control group, and based on Tillman’s work we suppose that fructose might have also increased metabolic rate thus maintaining body weight stable.

After short-term fructose feeding (4 weeks), C57BL/6 mice still presented neither increased adiposity nor adipocyte hypertrophy, although early signs of fat redistribution were found toward an increased visceral WAT. Long-term fructose feeding is supposed to stimulate adiposity, and Montgomery et al showed increased fat mass on visceral (perigonadal and retroperitoneal) and subcutaneous (inguinal) WAT depots after 8 weeks of fructose feeding (Montgomery, et al. 2015). Visceral and subcutaneous WAT have distinct functions, and traditionally visceral WAT has been associated with metabolic and cardiovascular disease risk (Wajchenberg, et al. 2002). For instance, visceral obesity can contribute to insulin resistance and coronary artery disease development in nonobese individuals (Filho, et al. 2006). However, Moreno-Indias et al showed in normal-weight subjects that macrophage-associated genes are upregulated in subcutaneous but not visceral WAT (Moreno-Indias, et al. 2016). It suggests that macrophages within subcutaneous fat may also contribute to the unhealthy phenotype seen in MONW individuals.

There is substantial evidence that dietary intake of high amount of fructose leads to the development of glucose intolerance, insulin resistance, and hepatic steatosis as reviewed elsewhere (Samuel 2011). Hypercholesterolemia was the only metabolic abnormality found by us after short-term fructose feeding (4 weeks). We believe that the time of fructose feeding required to develop further metabolic abnormalities such as insulin resistance needs to be longer than 4 weeks. Montgomery et al showed that long-term fructose feeding (8 weeks) did not change fasting glucose and insulin, but it promotes glucose intolerance and decreases plasma TG and non-esterified fatty acids (NEFA) in mice (Montgomery, et al. 2015). Tillman et al showed that glucose, triglycerides, and NEFA were not altered in C57BL/6 mice after 14 weeks of fructose feeding (Tillman, et al. 2014). Fructose stimulates gluconeogenesis, but it seems to produce only mild changes in blood glucose (Dirlewanger, et al. 2000). Although fructose does not increase insulin levels acutely because it does not induce pancreatic beta cell secretion of insulin like glucose, chronic exposure to fructose leads to hyperinsulinemia (Basciano, Federico, and Adeli 2005).

Insulin resistance includes impairment of fatty acid oxidation and utilization (Kelley and Goodpaster 2001), and hepatic TG content is a strong determinant of hepatic insulin resistance (Marchesini, et al. 1999). In the present study, short-term fructose intake increased liver weight despite no signs of hepatic steatosis, as pointed by TG content, TyG index and histopathological analysis (data not shown). Long-term studies have shown that fructose feeding promotes hepatic lipid accumulation of TG and DAG, but not ceramides (Montgomery, et al. 2015). The phenomenon is due to the upregulation of lipogenic pathways through the elevation of protein contents of both acetyl-CoA carboxylase isoforms (ACC1 and ACC2), fatty acid synthase (FAS) and stearoyl-CoA desaturase (SCD1) (Montgomery, et al. 2015).

Short-term fructose feeding did not change BAT mass, a result found by us and others (Montgomery, et al. 2015). We have previously shown in female C57BL/6 mice that BAT mass gain is a late event in BAT dysfunction during aging (Gonçalves, et al. 2017). We also showed in the present study that average LD size was increased by fructose, but cell size was not affected. We suppose that long-term fructose feeding would result in brown adipocyte hypertrophy, as well as white adipocyte hypertrophy. In aged male C57BL/6 mice, brown adipocyte hypertrophy is associated with BAT morphological abnormalities, lower thermogenesis, and glucose intolerance (Sellayah and Sikder 2014). LD hypertrophy is due to increased TG storage and, in humans, the TG content of thermogenic supraclavicular fat deposits may be an independent marker of whole-body insulin sensitivity, independent of BAT metabolic activation (Raiko, et al. 2015). In summary, these data show early modifications of BAT adipocyte architecture by fructose, that will likely lead to metabolic dysfunction in the future, but additional studies are necessary to confirm this hypothesis.

LD size regulation and lipid storage capacity are important to maintain normal biological functions, and its dysregulation results in the development of metabolic diseases such as obesity, diabetes, fatty liver and cardiovascular diseases. The nutritional response, hormones and environmental factors may also contribute to increased LD size and lipid storage (Beller, et al. 2010). LD growth occurs by the fusion of pre-existing LDs, lipid biosynthesis in situ or by lipid transfer from adjacent organelles, including endoplasmic reticulum. Although white and brown adipocytes accumulate large amounts of fat, LDs differ in size, number and protein content (Anand, et al. 2012). Brown adipocytes are filled with many relatively small sized LDs (multilocular) that are closely associated with mitochondria. This arrangement facilitates the accessibility of lipases to LD surface for the rapid release of fatty acids. LD biosynthesis and expansion are driven by complex and integrated mechanisms involving interactions with other organelles and enzymes for the expansion of the lipid core and the modulation of phospholipids monolayer composition (Barbosa, Savage, and Siniossoglou 2015).

The MONW phenotype is characterized by normal weight with increased adiposity and to investigate its underlying mechanisms is of ultimate importance. So far, MONW study in mouse models is limited, and fructose feeding is a potential model to investigate the mechanisms underlying the MONW phenotype. In C57BL/6 mice, it seems that fructose in the drinking water *ad libitum* can elicit an obesogenic response, but not fructose offered in the food. This very different response in mice to either fed fructose or given fructose liquid was not explored so far. Thus, further studies are necessary to determine the best route for fructose administration (chow or drink water). It is also necessary to investigate how long it takes to develop the MONW phenotype by fructose. Finally, it is important to know which cardiometabolic abnormalities commonly found in MONW humans fructose feed mice would develop (for instance, glucose intolerance, insulin resistance, low-grade inflammation, hypertriglyceridemia, and hypertension), and the time required to see these cardiometabolic changes.

## Conclusion

Short-term fructose feeding redistributes body fat, changes the proportion of BAT to visceral fat, and promotes BAT adverse remodeling. The novelty of the present work is that BAT displays early morphological signs of future metabolic dysfunction before cardiometabolic dysfunction itself, in a model of fructose-fed normal weight mice. A future direction is to determine the timing of cardiometabolic disturbance onset due to fructose overfeeding and BAT adverse remodeling in long-term fructose feeding in male and female C57BL/6 mice.

## Acknowledgements

Authors are thankful for Dilliane da Paixão Rodrigues Almeida for her technical assistance.

## Conflicts of interest

There are no conflicts of interest to declare.

## Funding source

This work was supported by the Foundation for Research Support of the State of Rio de Janeiro (FAPERJ, grant number E-26/210.525/2014) and by the Fluminense Federal University Pro-Rector of Research, Post-Graduate Studies and Innovation (PROPPI/UFF).

